# Generation of a mitochondrial protein compendium in *Dictyostelium discoideum*

**DOI:** 10.1101/2021.11.08.467494

**Authors:** Anna V Freitas, Jake T Herb, Miao Pan, Yong Cheng, Marjan Gucek, Tian Jin, Hong Xu

**Author notes:** Correspondence to: Hong Xu. These authors contributed equally to this work (Anna V Freitas, Jake T Herb).

## Abstract

The social amoeba *Dictyostelium discoideum* is a well-established model to study numerous cellular processes including cell motility, chemotaxis, and differentiation. As energy metabolism is involved in these processes, mitochondrial genetics and bioenergetics are of interest, though many features of *Dictyostelium* mitochondria differ from metazoans. A comprehensive inventory of mitochondrial proteins is critical to understanding mitochondrial processes and their involvement in various cellular pathways. Here, we utilized high-throughput multiplexed protein quantitation and homology analyses to generate a high-confidence mitochondrial protein compendium. Our proteomic approach, which utilizes quantitative mass spectrometry in combination with mathematical modeling, was validated through mitochondrial targeting sequence prediction and live-cell imaging. Our final compendium consists of 1082 proteins. Within our *D. discoideum* mitochondrial proteome, we identify many proteins that are not present in humans, yeasts, or the ancestral alpha-proteobacteria, which can serve as a foundation for future investigations into the unique mitochondria of *Dictyostelium*. Additionally, we leverage our compendium to highlight the complexity of metabolic reprogramming during starvation-induced development. Our compendium lays a foundation to investigate mitochondrial processes that are unique in protists, as well as for future studies to understand the functions of conserved mitochondrial proteins in health and diseases using *D. discoideum* as the model.

## Introduction

*Dictyostelium discoideum*, a social amoeba, is a well-established model organism to study eukaryotic cellular processes such as cell motility, chemotaxis, and differentiation (Bozzaro, 2013). Under normal nutrient conditions, *D. discoideum* grows axenically through binary fission (Kessin, 2001). However, upon starvation, amoebae secrete cAMP, which attracts neighboring cells to aggregate together and form a multicellular mound. Cells in a mound move collectively as a slug toward light, heat, or humidity to find a suitable environment. The slug eventually matures into a fruiting body consisting of two major types of differentiated cells, spore cells that will start a new life cycle and stalk cells that form a stalk to hold the spore aloft (Kay, 1982). As many of the aforementioned biological processes are intertwined with cellular energetics, investigation of mitochondrial biogenesis and functions is an emerging area in *D. discoideum* research (Francione *et al*., 2011; Pearce *et al*., 2019).

The *D. discoideum* mitochondrial genome is ∼56 kb, circular, double-stranded DNA that encodes two ribosomal RNAs, 18 transfer RNAs (tRNAs), five open reading frames without annotated function, and 38 proteins including 18 subunits of the electron transport chain complexes and 15 ribosomal proteins (Ogawa *et al*., 2000). Phylogenetic studies reveal that Amoebazoa diverged before Opisthokonta, but after the divergence of Plantae (Baldauf and Doolittle, 1997), and are more closely related to animals than plants. Notably, the *Dictyostelium* mitochondrial genetic system possesses a few differences from metazoans (Pearce *et al*., 2019). *D. discoideum* mitochondrial DNA (mtDNA) has four introns in *cox1/2* genes and utilizes universal codons (Angata *et al*., 1995; Ogawa *et al*., 2000), a common feature of most plants’ mitochondria (Jukes and Osawa, 1990; Cho *et al*., 1998). The universal genetic code and the lack of a full set of tRNA genes on the *Dictyostelium* mitochondrial genome indicate that some nuclear-encoded tRNAs are likely imported into mitochondria to support the organellar translation. Additionally, the electron transport chain in *Dictyostelium* contain an additional component compared to its metazoan counterparts: an alternative oxidase (AOX) (Pearce *et al*., 2019), which is found across eukaryotic clades besides animals (McDonald *et al*., 2008). AOX is highly expressed during vegetative growth, but its expression level is markedly reduced upon starvation, suggesting a potential metabolic reprogramming occurs during starvation-induced development (Jarmuszkiewicz *et al*., 2002). Interestingly, either reduction of mtDNA content or disruption of the *rps4* locus (encoding mt-ribosomal protein S4) on mtDNA impairs aggregation and slug phototaxis but has no impact on vegetative growth (Chida, 2004; Chida *et al*., 2008), suggesting that mtDNA, and most likely an intact oxidative phosphorylation system is essential to initiate the development program. On the other hand, pharmacological inhibitions of either Complex I or Complex V can induce aggregation, even though mitochondrial respiration appears to increase at the beginning of starvation (Kelly *et al*., 2021). Therefore, the interplay betweesn mitochondrial function and *Dictyostelium* development remains to be explored.

Despite the growing interest in using *D. discoideum* as a model organism to study many conserved mitochondrial processes and some unique biology, a comprehensive list of the mitochondrial proteins has yet to be established. A recent proteomic study detected 294 proteins in *D. discoideum* mitochondria (Mazur *et al*., 2021), which we believe is far from complete. Nuclear-encoded mitochondrial proteins, which constitute over 90% of the total mitochondrial proteome, are synthesized in the cytoplasm and then imported to mitochondria. It is estimated that the import of ∼60% of these proteins relies on a positively charged, N-terminal mitochondrial targeting sequence (MTS) (Vögtle *et al*., 2009). Computational approaches that integrate machine learning and known biological data are frequently used to predict mitochondrial targeting based on the presence of an MTS (Almagro Armenteros *et al*., 2019). However, this method is insufficient to capture all mitochondrial proteins, as most proteins on the outer membrane and in the inner membrane space, and some inner membrane proteins rely on alternate translocation mechanisms. An alternative computational approach leverages sequence homology to known mitochondrial protein compendiums that were generated using mass spectrometry (MS)-based proteomic discovery (Pagliarini *et al*., 2008; Morgenstern *et al*., 2017). However, to compensate for the rapid evolution of the mitochondrial genome, nuclear-encoded mitochondrial proteins evolve faster than other nuclear-encoded proteins (Cole *et al*., 1995; Sloan *et al*., 2014; Havird *et al*., 2015; Li *et al*., 2017; Yan *et al*., 2019). Thus, some *Dictyostelium* mitochondrial proteins may escape the homology search, and protist-specific mitochondrial proteins will certainly be missed.

In this study, we combined quantitative proteomics and mathematical modeling to identify over 900 high-confidence mitochondrial proteins, which were validated through both *in silico* and fluorescent microscopy analyses. We further complemented the proteomics-based mitochondrial protein discovery with bioinformatic approaches to create a compendium of 1082 *D. discoideum* mitochondrial proteins. We also discuss conserved *D. discoideum* mitochondrial proteins that may be used as the basis of validating mitochondrial proteins in other organisms, as well as unique features of the mitochondrial proteome in *D. discoideum*.

## Results and Discussion

### Mitochondrial protein discovery using quantitative proteomics

To identify putative *D. discoideum* mitochondrial proteins, we searched for *Dictyostelium* homologs of 1136 human mitochondrial proteins listed in the Human MitoCarta 3.0 (Morgenstern *et al*., 2017; Rath *et al*., 2021), and retrieved 616 proteins (Figure 1A, Table S1). This number is much less than known mitochondrial proteins in humans (1136) and baker’s yeast (901) (Pagliarini *et al*., 2008; Rath *et al*., 2021). We posited that mitochondrial proteome might be highly divergent between *D. discoideum* and humans, and many *Dictyostelium* mitochondrial proteins might be missed from this bioinformatic curation. We, therefore, took a proteomic approach to directly identify mitochondrial proteins in *D. discoideum* (Figure 1A). From AX2 axenic cultures, we prepared mitochondria isolates—both crude and highly purified—through Percoll gradient ultracentrifugation. We performed tandem mass tag (TMT) liquid chromatography-mass spectrometry (LC-MS) on both mitochondria isolates and included AX2 whole-cell lysate as the control. A total of 6,892 proteins were captured in all samples (Table S2).

**Figure 1.**
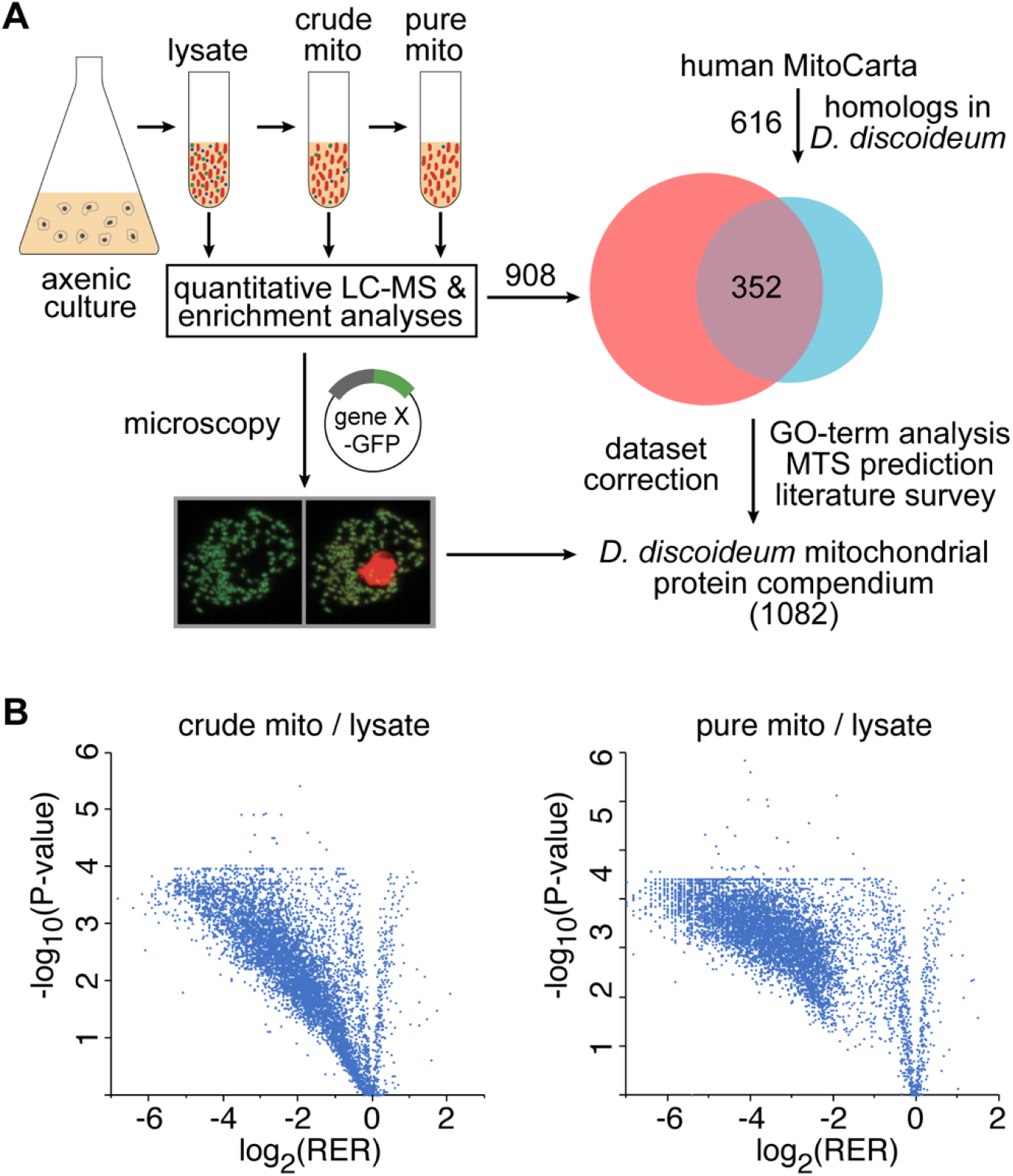
Curation of a comprehensive mitochondrial proteome in *Dictyostelium discoideum*. (A) A homology search against human mitochondrial protein sequences yielded a list of 616 putative *D. discoideum* mitochondrial proteins. To identify additional mitochondrial proteins, we performed quantitative MS for proteins identified in the whole-cell lysate, as well as in crude and purified mitochondrial samples. Microscopy was used to validate MS and enrichment analyses. After correcting the proteomic dataset based on microscopy results, we incorporated the homology and proteomic analyses to yield a comprehensive mitochondrial protein compendium. (B) Volcano plots displaying the −log10 (p-value) versus log2 (relative enrichment ratios) of proteins in crude mitochondrial or pure mitochondrial versus whole-cell lysate samples. Data are presented as means (n = 3).

A limitation of identifying organellar proteins from their subcellular fractions alone is that high-abundance contaminants are often co-purified and result in false-positive hits. To address this issue, we assessed the probability of a protein localizing to mitochondria by comparing its relative enrichment in mitochondrial preparations to a list of 47 authentic mitochondrial proteins that includes components of electron transport chain complexes and conserved enzymes in citrate cycles (Table S3). We first calculated the ratio of a protein’s abundance in the mitochondria isolates, both crude and highly purified, versus its abundance in the whole-cell lysate. The resulting value, indicating its enrichment in mitochondrial preparations, was further normalized to the average enrichment ratio of the 47 reference mitochondrial proteins, to compute the relative enrichment ratio (RER). Overall, a protein’s RER in crude mitochondria isolate is in accordance with that in purified mitochondria (Figure 2A). However, the distribution of RERs appears continuous in crude mitochondria (Figure 1C), but clusters into two distinct populations in purified mitochondria (Figure 1C), which allowed us to determine a proper threshold of RER for mitochondrial proteins using mathematical modeling. Thus, we proceeded to analyze the RER for purified mitochondria only.

**Figure 2.**
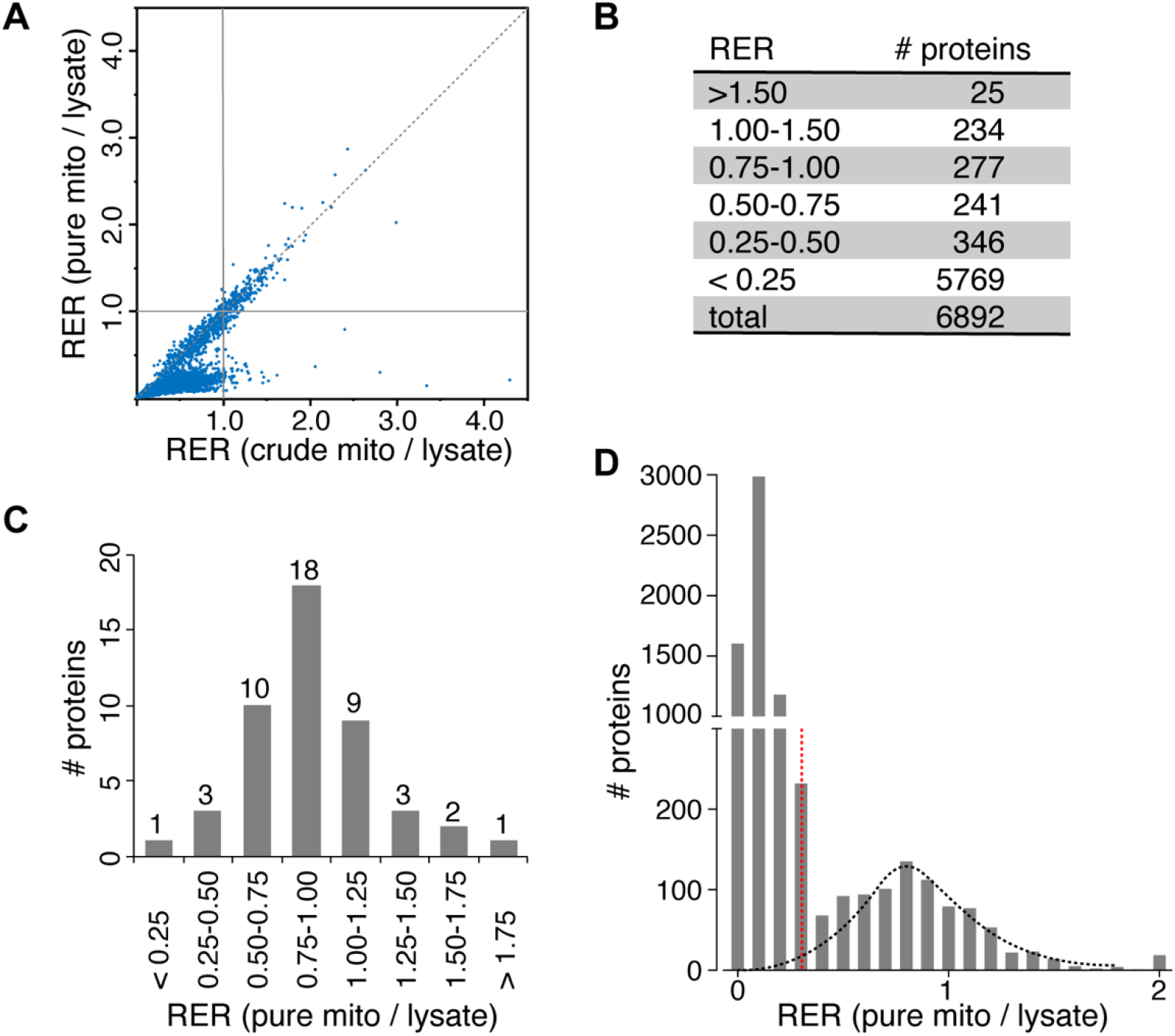
Prediction of mitochondrial localization based on relative enrichment analysis. (A) Crude mitochondrial RER versus pure mitochondrial RER for all samples quantified in the proteomic discovery experiment. The dashed line represents equal crude and pure RERs. Data are presented as means (n = 3). (B) Summary table of pure mitochondrial relative enrichment ratios for all proteins in the protein discovery experiment. (C) Distribution of core *D. discoideum* mitochondrial ETC, OXPHOS, and TCA cycle proteins (n = 47) based on RER. (D) The distribution of mitochondrial (n = 908) and non-mitochondrial proteins (n = 5984) based on RER follow a Gaussian mixture model, in which a cutoff value (red, RER = 0.343) separates two normal curves.

In principle, a true mitochondrial protein would be co-purified with the reference mitochondrial proteins in pure mitochondrial isolates, and its RER should be 1.0. However, the RER distribution of these 47 reference proteins (Figure 2B) appears as a normal curve centered around 1.0, suggesting that many mitochondrial proteins may have an RER below 1.0. Among all proteins profiled using TMT-based LC-MS, only 259 have an RER higher than 1.0 in purified mitochondria (Figure 2C). We posit that different mitochondria proteins might be degraded to different extents, based on their intrinsic stability, during the procedure of mitochondrial purification, which involves overnight ultracentrifugation. Therefore, it is necessary to determine a proper RER value to differentiate mitochondrial proteins from non-mitochondrial proteins. We applied the expectation-maximization (EM) approach to a gaussian mixture model (GMM) to bin all proteins into two clusters: non-mitochondrial and mitochondrial proteins based on their RER values (Figure2D). We chose an RER cutoff of 0.343 (Figure 2D) and assigned a total of 908 proteins having an RER higher than 0.343 as putative mitochondrial proteins (Table S4). GMM predicts that less than 17% proteins in the mitochondrial cluster, and only 0.1% proteins in the non-mitochondrial cluster would spill over to the other group (Figure 2D), which corresponds to an 83% recovery rate and 7% false discovery rate, respectively.

### Validation of mitochondrial protein discovery based on quantitative proteomics

To validate the accuracy of RER-based mitochondrial protein discovery, we first assessed the recovery rate of putative mitochondrial proteins *in silico*. Many mitochondrial proteins possess an N-terminus mitochondrial targeting sequence (MTS) that directs the import of nuclear-encoded mitochondrial proteins into the mitochondrial matrix (Backers, 2017). Overall, 24% of all proteins retrieved in the proteomics discovery experiment contain an MTS (Table S4). Importantly, 94% of these MTS-bearing proteins had an RER greater than 0.343 (Figure 3A). On the contrary, 96% of proteins that were destined to other organelles such as the ER, Golgi, lysosomes, vacuoles, or secretory pathway had an RER less than 0.343. These analyses demonstrate that a cutoff value of an RER at 0.343 effectively separates mitochondrial proteins from non-mitochondrial proteins.

**Figure 3.**
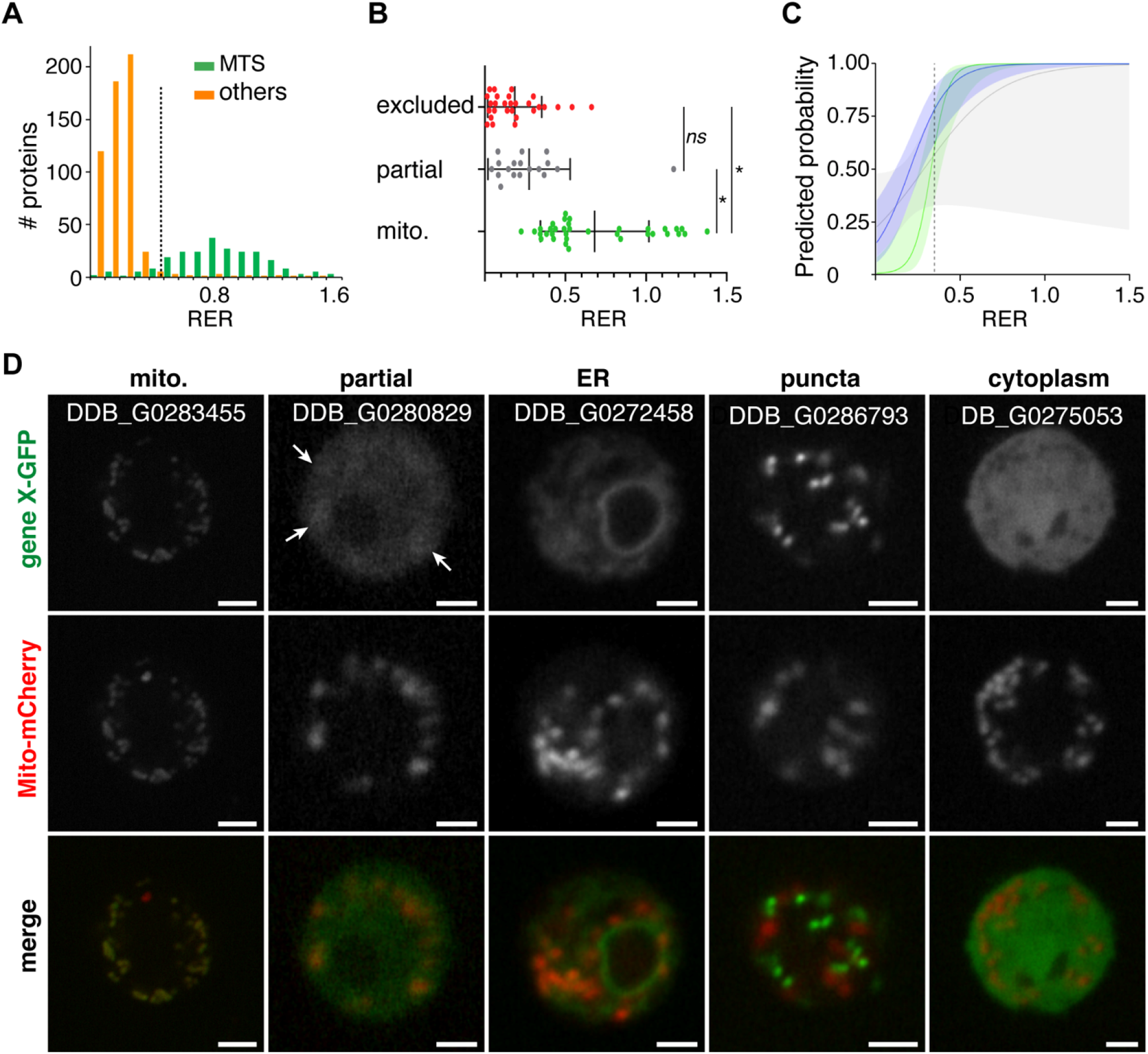
Validation of proteomic discovery and enrichment analyses. (A) Distribution of all proteins quantified with a predicted mitochondrial targeting sequence (MTS) or signal peptide sequence. The dashed line represents the RER cutoff for predicting mitochondrial localization (0.343). (B) Dot plot with interquartile ranges of proteins that had mitochondrial localization (green), partial mitochondrial localization (gray), or were excluded from the mitochondria (red). *, *p* ≤ 5 × 10^−5^. (C) Predicted probabilities and confidence intervals as analyzed by logistic regression for mitochondrial (green), partial mitochondrial (gray), and either mitochondrial or partial (blue) localization. The dashed line represents the RER cutoff for predicting mitochondrial localization (0.343). (D) Representative live-cell confocal images of axenic stage *D. discoideum* expressing GFP-tagged genes of interest leveled with cytochrome oxidase c subunit IV tagged with mCherry (Mito-mCherry). Arrowheads denote areas of partial localization.

We also surveyed the localization of 81 proteins recovered in LC-MS (Table S5), using fluorescent microscopy. These proteins were selected on the basis that their subcellular localization has not been annotated previously as mitochondrial, and their RERs are randomly distributed from 0.1 to 1.5. Each protein was tagged with GFP at its C-terminus and co-expressed with an MTS-mCherry fusion protein, which marks mitochondria in *D. discoideum* AX2 cells. Among the 81 proteins, 90% of proteins with an RER higher than 0.343 showed complete or partial mitochondrial localization (Figure 3B, 3D), whereas only 5% of proteins with an RER less than 0.343 showed mitochondrial localization (Figure 3B, 3D), demonstrating a strong positive correlation between RER value and probability of mitochondrial localization (Figure 3B). Moreover, logistic regression analysis on the localization pattern of these 81 proteins predicts that a protein has more than a 78% probability of localizing to the mitochondria if its RER is higher than 0.343 (Figure 3C).

### A comprehensive mitochondrial protein compendium in *D. discoideum*

To further improve the coverage and accuracy of the mitochondrial protein discovery, we revised the list based on the *in vivo* microscopy validation by removing four non-mitochondrial localizing proteins and adding two mitochondrial localizing proteins. We also integrated three sets of mitochondrial protein discovery: the aforementioned list of mitochondrial proteins identified from quantitative proteomics analyses, those retrieved from homology detection, and those retrieved during a gene ontology search for mitochondrial genes. Among the 616 *D. discoideum* homologs of human mitochondrial proteins (Table S1), 352 proteins have an RER higher than 0.343 and hence were already included in the list, 223 proteins have an RER lower than 0.343, and 41 proteins were not captured in LC-MS. Among the 264 proteins that were not included in the list, 113 proteins do not have a predicted MTS (Table S1), whereas their human homologs have MTSs, suggesting these proteins might localize to other cellular compartments in *D. discoideum*. An exception is ribosomal protein S14 (O21035), which is encoded in the nuclear genome in humans but is encoded in the mitochondrial genome in *D. discoideum*, and thus contains an MTS in human cells but lacks one in *D. discoideum*. We added those remaining 152 proteins to the list, as well as 32 proteins with mitochondrial gene ontologies that had not emerged during the proteomic or homology analysis. The final compendium consists of 1082 high-confidence mitochondrial proteins in *D. discoideum* (Table S6).

### Characterization of the *D. discoideum* mitochondrial proteome

Out of the 1082 *D. discoideum* mitochondrial proteins, there are 627 and 458 proteins that have homologs in the mitochondrial proteome of human and *Saccharomyces cerevisiae*, respectively (Figure 4A), indicating that *D. discoideum* mitochondria are more closely related to mitochondria in metazoans than fungi. Only 324 *D. discoideum* mitochondrial proteins have homologs in *Rickettsia prowazekii* (Figure 4A), an α-proteobacteria that is closely related to the mitochondrial ancestor. Overall, a total of 313 proteins, representing 28.9% of the *D. discoideum* mitochondrial proteome, have no homologs in the whole proteome of humans, *S. cerevisiae* or *R. prowazekii* (Figure 4A, Table S6), indicating that a large fraction of *D. discoideum* mitochondrial proteins was evolved *de novo* after the divergence of Amoebozoa. Moreover, 75 *D. discoideum* mitochondrial proteins (6.9%) have no homologs in *D. purpureum* (Figure 4A), a closely related species of social amoeba, further substantiating the fast-evolving nature of the amoeba mitochondrial proteome.

**Figure 4.**
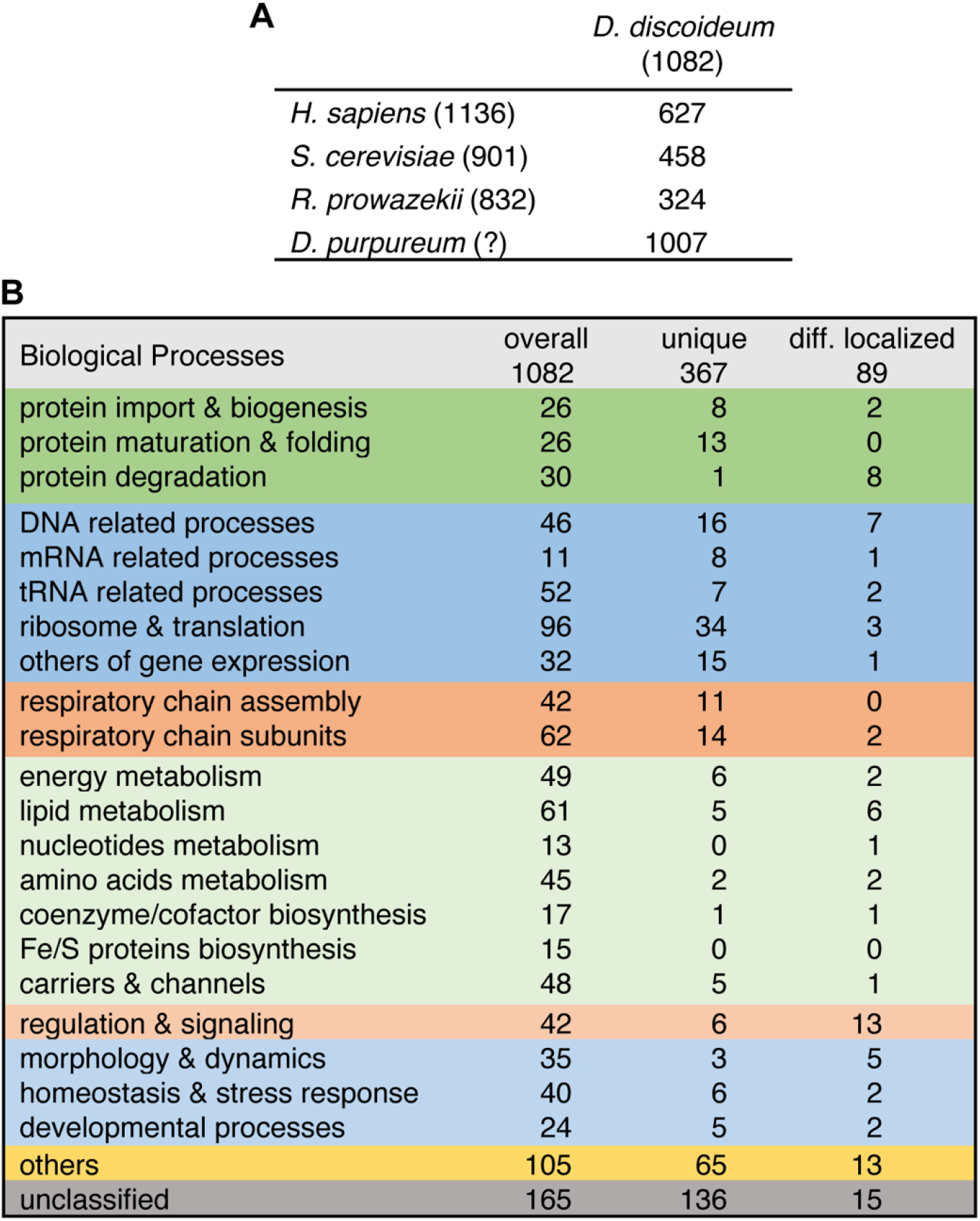
Categorization of the *D. discoideum* mitochondrial protein compendium. (A) *D. discoideum* mitochondrial proteins with homologs in *H. sapiens* or *S. cerevisiae* mitochondrial proteomes, or the complete proteomes of the alphaproteobacteria *R. prowazekii* or *D. purpureum*. (B) The functional categorization of all *D. discoideum* mitochondrial proteins (overall, n = 1082), those that lack human homologs (unique, n = 367), and those that have human homologs, but homologs were not annotated as mitochondrial proteins (diff. localized, n = 89).

There are 89 *D. discoideum* mitochondrial proteins (8.2%) with human homologs that had not been annotated as mitochondrial proteins (Figure 4B, Table S6). Among these 89 proteins, 74 were also not annotated as mitochondrial proteins in yeast, including 32 that had homologs in *S. cerevisiae*. Given the estimated false-discovery rate of our compendium, the localization of these proteins needs to be experimentally accessed. Nonetheless, there are a few examples, such as the RNB domain-containing protein (DDB_G0288469) and tRNA-binding domain-containing protein (DDB_G0349377), both of which have predicted MTSs and are likely targeted to the mitochondrial matrix. The human homologs of DDB_G0288469, DIS3-like exonuclease 2, and DDB_G0349377, rhomboid-related protein 4, were not included in the human compendium (Rath *et al*., 2021), despite evidence that the yeast homolog of DIS3-like exonuclease 2 localizes to the mitochondria (Pagliarini *et al*., 2008), and that rhomboid-related protein 4 has been partially shown to localize to the mitochondria. The mitochondrial localization of their *D. discoideum* homologs substantiates these two proteins might indeed localize to the mitochondria and indicates that our compendium can complement previous studies toward a more comprehensive discovery of mitochondrial proteins in other organisms.

Additionally, we categorized the *D. discoideum* mitochondrial proteome using PANTHER biological function or protein family classifications (Figure 4B, Table S6). Proteins involved in mitochondrial gene expression and metabolism comprise the largest fractions of all mitochondrial proteins, over 20% for each category. Other proteins are involved in mitochondrial protein homeostasis, the electron transport chain, redox signaling and metabolism, and regulation of mitochondrial morphology and dynamics. A large fraction of *D. discoideum* mitochondrial proteins, approximately 15%, have no classified functions based on PANTHER analyses (Figure 4B).

### *D. discoideum*-specific mitochondrial proteins

Proteins involved in gene expression consisted of a large fraction of *D. discoideum* specific mitochondrial proteome (Figure 4B), reflecting that the *D. discoideum* mitochondrial genome is more complex than human mtDNA. On the contrary, few metabolism proteins emerged in the list (Figure 4B), suggesting that metabolic processes are highly conserved between *D. discoideum* and metazoans. Here, we expand upon a few of the unique features of the *D. discoideum* mitochondrial protein compendium.

#### Mosaic nature of mitochondrial ribosomes

Mitochondrial ribosomes (mitoribosomes), ribosomal assembly factors, and other proteins involved in translation represented 8.9% and 9.3% of the overall and unique mitochondrial protein compendium, respectively. While mitoribosomes are thought to be evolved from bacterial ribosomes, these two differ greatly with regards to their structure, function, as well as their composition of proteins and RNAs. We identified 51 proteins that are predicted to be mitoribosomal proteins, including 13 proteins that did not share significant homology with any *H. sapiens, S. cerevisiae*, or *R. prowazekii* proteins (Table S7). Interestingly, *D. discoideum* mitoribosomal proteins belong to families across several taxonomic groups: 35 proteins belong to mammalian mitoribosomal protein families (28s and 39s), 2 belong to eukaryotic cytosolic ribosomal protein families (60s), 9 belong to yeast mitoribosomal protein families (37s and 54s), 2 belong to chloroplast or bacterial ribosomal protein families (30s and 50s), 1 is from archaea, and 2 are universally conserved among prokaryotes and eukaryotes. It has previously been shown that cytosolic ribosomes tether to the mitochondrial outer membrane. Hence, the recovery of the 60S ribosomal protein L22 could be the result of the association of cytoplasmic ribosome with the mitochondrial outer membrane rather than its localization in the matrix (Gold *et al*., 2017). Nonetheless, the presence of proteins representing multiple mitoribosome lineages suggests that there may be *D. discoideum* or protist-specific mechanisms to process mitochondrial transcripts and to regulate mitochondrial translation. Further validation of these findings is necessary as the composition and structure of the *D. discoideum* mitoribosome have yet to be resolved.

#### Mitochondrial DNA and RNA processing factors

Among the list of unique proteins are 24 candidate mtDNA and mtRNA processing factors including five endonucleases and a pentatricopeptide repeat (PPR)-containing protein A (PtcA). Bioinformatic analysis suggests that PtcA belongs to the mitochondrial group I intron splicing family. PPR proteins, defined by tandem PPR domains, are implicated in several different mitochondrial gene expression processes including translation initiation, and ribosomal stabilization (Manna, 2015). The number of PPR proteins that are encoded in an organism varies greatly: terrestrial plants, such as *Arabidopsis thaliana*, have upwards of 450 PPR proteins, while humans have 7 (Lurin *et al*., 2004; Lightowlers and Chrzanowska-Lightowlers, 2013). *D. discoideum* has 12 PPR-domain containing peptides, including PtcA, reflecting a greater complexity of *D. discoideum’s* mitochondrial genome compared to that of metazoans (Manna *et al*., 2013).

#### Divergent evolution path of lipid biosynthesis

The mevalonate pathway, which produces five-carbon blocks for the synthesis of diverse biomolecules such as cholesterol and coenzyme Q10, is an essential and highly conserved process in eukaryotes, archaea, and some bacteria. In animals and fungi, the mevalonate pathway takes place in ER, and 3-hydroxy-3-methylglutaryl (HMG)-coenzyme A (CoA) reductase (HMGR), a key enzyme in this pathway that converts HMG-CoA to mevalonate, localizes in the ER and peroxisomes (Chin *et al*., 1984; Keller *et al*., 1986; Burg and Espenshade, 2011). HMGR2, one of two HMG reductases in *D. discoideum*, is recovered in our compendium and contains a predicted MTS, suggesting that it likely localizes to the mitochondrial matrix. Additionally, HGSA, one of the two HMG-CoA synthases, also emerged as a mitochondrial protein. Our mitochondrial protein discovery suggests that mevalonate metabolism may take place in mitochondria in *D. discoideum*, highlighting the evolutionary divergence of some metabolic pathways that originated from the common mitochondrial ancestor.

### Implication of mitochondrial function in multicellular development

*D. discoideum* with reduced mtDNA or a disruption of the gene encoding mt-ribosomal protein S4 display no defect in vegetative growth but have impaired starvation-induced development, suggesting that mitochondrial respiration is necessary for multicellularity (Chida, 2004; Chida *et al*., 2008). However, contrasting evidence has demonstrated a significant decrease in mitochondrial respiration after respiration, and accordingly, a decreasing expression of many respiration complexes (Kelly *et al*., 2021). To understand potential regulations of mitochondrial function in multicellular development, we retrieved RNA sequencing data using the *Dictyostelium* gene expression database, dictyExpress (Parikh *et al*., 2010; Stajdohar *et al*., 2017).

Overall, there was a decrease in the expression of mitochondrial genes within our compendium over the 24-hr development time course (Figure 5A). A similar pattern is observed in proteins that are involved in mitochondrial DNA maintenance and gene expression. Interestingly, despite the decrease in gene expression machinery (Figure 5B), over half of the mitochondria-encoded genes in the dataset (19 of 35) were upregulated (log2FC ≥ 1) after starvation induction (Figure 5C). Further, in examining all respiratory chain complexes, 12 nuclear-encoded ETC subunits had a higher expression level (log2FC ≥ 1) at or after 12 hours of starvation (Figure 5D), besides the 10 nuclear or mitochondrial-encoded subunits that show a burst of expression in the first 4 hours after the starvation (Figure 5D). The complex pattern of mitochondrial gene expression, particularly the upregulation of electron transport chain complex subunits during the development suggests potential roles of mitochondrial respiration in *Dictyostelium* development, and that both nuclear and mitochondrial-encoded proteins are likely implicated in these processes.

**Figure 5.**
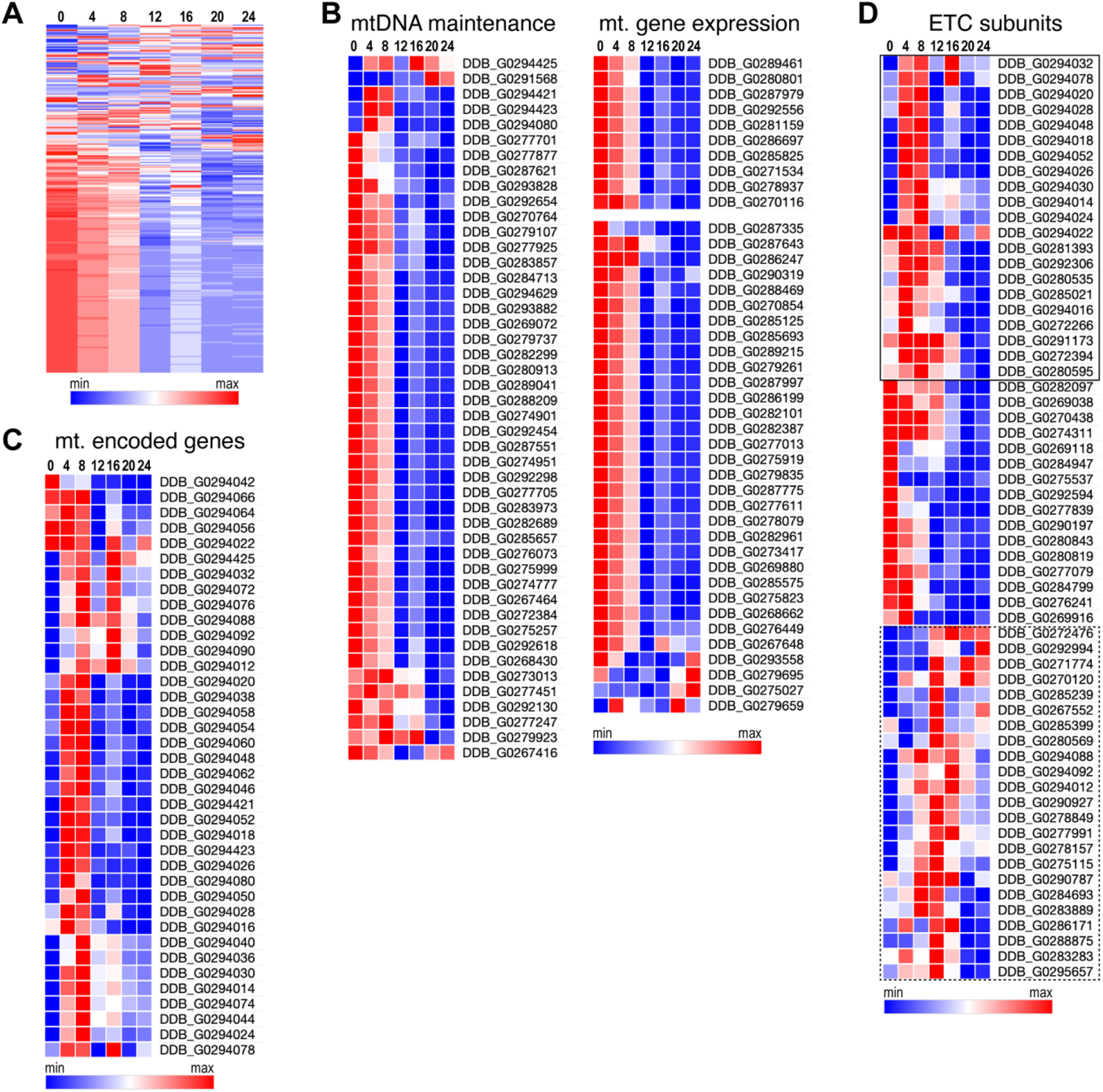
Expression profile of mitochondrial genes during *D. discoideum* development. Heatmaps showing normalized RNA-seq analysis values of (A) all mitochondrial proteins within the compendium and (B-D) specific mitochondrial proteins during the 24-hour starvation-induced development cycle. Each column of heat maps represents the time in hours after developmental induction. Rows are clustered by similarity in gene expression profiles. Boxes outline ETC subunits showing increased expression 4 hours after starvation (solid) and 12 hours after starvation (dashed).

## Conclusion

Here, we generated the most comprehensive list of mitochondrial proteins in *D. discoideum* to date. Our compendium lays the foundation for future studies to understand the functions of conserved mitochondrial proteins in health and diseases using *D. discoideum* as the model. It also provides an entry to study many fascinating mitochondrial processes that are unique in protists. Additionally, thorough comparative genomics, our compendium will complement mitochondrial protein discovery in other organisms and may shed light on the evolution of mitochondrial proteome and processes.

## Materials and Methods

### Cell culture and transformation

*Dictyostelium discoideum* AX2 cultures were maintained in HL5 medium at 22 °C (Fey *et al*., 2007). Transformants were generated via electroporation as previously described (Gaudet *et al*., 2007), with modifications. After electroporation (BioRad Genepulser), cells were incubated on ice for 10 minutes. Subsequently, cells were transferred from the cuvette with 2 mL of HL5 and plated onto 12-well tissue culture plates. After 24 hours, transformants were selected with Genectin and/or Blasticidin S (Thermo Fisher, 10 μg/mL each) in HL5.

### Protein mass spectrometry

#### Mitochondrial isolation

Cells were harvested at a concentration of 1-3 × 10^6^ cells/mL and resuspended at 2 × 10^7^ cells/mL in 800 μL of Reagent A from the Mitochondrial Isolation Kit for Cultured Cells (Thermo Fisher #89874) on ice. Cell lysis and crude mitochondrial preparation were performed as previously described with modifications (Graham, 1999; Glancy and Balaban, 2011). Cells were lysed with 35 strokes of a Dounce homogenizer followed by the addition of an equal volume of Reagent C. Whole-cell lysate samples were stored at -80 °C or were centrifuged three times (700 x g, 10 minutes, 4 °C) to purify the mitochondria. For each centrifugation step, the supernatant was transferred to a fresh 1.5 mL tube. The crude mitochondrial lysate was used immediately for purification or was stored at -80 °C.

To generate purified mitochondrial isolates, Percoll gradient centrifugation was performed as follows. Lysis suspension (1-2 mL) was added to the top of a Percoll (Cytiva) and Development Buffer (DB) (5 mM Na_2_HPO_4_, 5 mM KH_2_PO_4_, 1 mM CaCl_2_, 2 mM MgCl_2_, pH = 6.5) solution (8 mL, 30% Percoll) in a 10 mL ultracentrifuge tube. Ultracentrifugation (68,000 x g, 40 minutes) yielded three distinct layers. The top layer, containing contaminants, was discarded. The middle layer, containing mitochondria, was transferred into a fresh 2 mL tube. Aliquots of the mitochondrial suspension were topped off with 500 μL of DB, then centrifuged (13,000 x g, 10 minutes, 4 °C). Following centrifugation, the supernatant was aspirated, and the mitochondria-containing pellet was maintained on ice. To lyse the mitochondria, the pellet was washed with 2 mL of DB, centrifuged (10,000 x g, 10 minutes, 4 °C), and resuspended in 2 mL of DB with urea (8M). Protein yield was quantified via Bradford Assay (BioRad) according to the manufacturer’s protocol.

#### Relative protein quantification

Resuspended cell pellets were lysed via pulsed sonication, then sequentially reduced, alkylated, and digested overnight with trypsin. The protein digests were labeled with 10-plex Tandem Mass Tag (TMT) reagents (Thermo Fisher Scientific) (Dayon *et al*., 2008), then were pooled and desalted. To separate the peptide mixtures into 24 fractions, high pH reversed-phase liquid chromatography was performed (Yang *et al*., 2012). Each fraction was analyzed on an Orbitrap Lumos (Thermo Fisher Scientific) nanoLCMS system.

Peptide and proteins were identified as described in He *et al*., (2020). In brief, the resulting LCMS raw data were searched against a database downloaded from dictybase.org using the Sequest HT algorithm on the Proteome Discoverer 2.4 platform (Thermo Fisher Scientific). Three groups of samples were normalized to 47 reference mitochondrial proteins.

The relative enrichment (RE) was defined as the ratio of a protein’s enrichment from a crude or purified mitochondria sample over its enrichment from a whole-cell lysate sample. To calculate the relative enrichment ratio (RER), REs were normalized such that the average RER of 47 known mitochondrial (TCA cycle, ETC, or OXPHOS) proteins is 1 (Table S3). The RER presented is a median value of three biological replicates.

### Mathematical modeling

To classify proteins as mitochondrial or non-mitochondrial based on their RER, the RER distribution of isolated proteins was fit to a Gaussian mixture model (GMM) using the Expectation-Maximization (EM) algorithm in R (Benaglia *et al*., 2009). The cutoff value (RER = 0.343) was four times the standard deviation plus the mean of curve 1 (representing non-mitochondrial proteins), such that 99.9% of the proteins below the cutoff were contained within curve 1.

### Bioinformatic analyses

#### Homology analyses

Protein sequence homology was established by BlastP expect < 0.001 and bit-score > 40 (Pearson, 2013), or by HMMER sequence e-value < 0.01. Subcellular localization was predicted using TargetP-2.0 (Almagro Armenteros *et al*., 2019). Biological functions for all proteins in the *D. discoideum* and human mitochondrial proteome were manually categorized from biological function or protein family classifications provided from PANTHER (The Gene Ontology Consortium *et al*., 2021).

#### *In silico* dataset correction

Two strategies were implemented to supplement the mitochondrial protein discovery. The top *D. discoideum* homolog of human mitochondrial proteins (Table S1, Rath *et al*., 2021) were curated. Additionally, *D. discoideum* proteins annotated with the gene ontology term “mitochond*” on AmiGO were selected (Carbon *et al*., 2009). Proteins within these lists were integrated into the final mitochondrial compendium so long as they had a predicted mitochondrial targeting sequence if their human homolog also had a predicted mitochondrial targeting sequence, except in cases where the *D. discoideum* protein was mitochondrial-encoded.

#### RNA sequencing data visualization

Normalized RNA-seq data from Parikh et al. (2010) was retrieved using dictyExpress (Stajdohar *et al*., 2017). For the 1082 proteins in the mitochondrial compendium, only 1075 corresponding genes were present in the dataset. To compare the gene profiles, data were scaled to a mean of 0 and standard deviation of 1 using the scale function in R. For the overall mitochondrial expression profile, genes and timepoints were ordered using hierarchical clustering (heatmaps.2). For the profiles of individual biological processes, scaled data (Table S8) were imported into the matrix visualization software Morpheus (https://software.broadinstitute.org/morpheus) and ordered via hierarchical clustering with one minus Pearson’s correlation as the distance metric and average as the linkage method. Gene upregulation was determined by a log2 fold-change ≥ 1.

### Library generation for imaging verification

To evaluate the efficacy of our mitochondrial protein identification, 98 proteins were selected to be GFP tagged so that their localization could be assessed via fluorescence microscopy (Table S5). None of the proteins selected had a gene ontology annotation that indicated mitochondrial localization. Four proteins selected for verification had homologs listed in the human mitochondrial proteome.

All genes were synthesized by Gene Universal. Of the 98 genes submitted for synthesis, 85 were generated as inserts in pDM323, a *D. discoideum* extrachromosomal expression vector with G418 resistance and a C-terminal GFP tag (Veltman *et al*., 2009); 6 genes were generated as inserts in the shuttle vector puC57 and were subsequently cloned into pDM323 between BglII and SpeI sites using the In-Fusion® HD Cloning Kit (Takara Bio USA) and confirmed by sequencing. The other 7 genes were unable to be synthesized, such that only 92 proteins were screened. Of these 92 genes, only 81 were successfully expressed in *D. discoideum*. The RER of the proteins that were verified were as follows: 14 proteins with a RER > .75, 11 proteins with a RER = 0.75-0.5, 24 proteins with a RER = .5-0.25, and 32 proteins with RER < 0.25. To observe mitochondrial localization, the mitochondrial targeting sequence of respiratory cytochrome oxidase c subunit IV fused with mCherry (CoxIV-mCherry) was cloned into pDM326, a *D. discoideum* extrachromosomal expression vector with Blasticidin S resistance (Veltman *et al*., 2009). All primers for cloning are listed in Table S9.

### Live-cell imaging

To image cells in the axenic phase, cells (200 μl) were transferred to 8-well glass chambers 7 to 10 days post-transformation. Cells were allowed to adhere to the bottom of the chamber for 30 minutes before the media was aspirated. The media was replaced with 1x PBS after three washes (200 μl for all). Confocal images were collected on a PerkinElmer Ultraview system (Zeiss Plan-apochromat 63x/1.4 oil lens, Volocity acquisition software, Hamamatsu Digital Camera C10600 ORCA-R2, Immersol immersion oil). Images (0.5 μm z-step) were analyzed with ImageJ (National Institutes of Health) and formatted in Adobe Photoshop.

### Code and data availability

Proteomics data are deposited at ProteomXchange (PXD029101). R code is available from Anna Freitas’s GitHub repo https://github.com/freitasav/DD-mitoproteome.

### Quantification and Statistical Analyses

All data were presented as the mean ± SD unless otherwise indicated. P values were calculated in R using a one-way analysis of variance (ANOVA) followed by Tukey’s post-hoc test to test for the effect of RER on mitochondrial localization. Statistical significance of difference was considered when p < 0.05.

To predict the probability of localization based on RER, outliers were identified and removed from the microscopy validation dataset based on the interquartile method (median + 1.5 SD). Data were analyzed using a binomial logistic regression (glm function in R) with *excluded from the mitochondria* as the reference level, and *partial mitochondrial localization, mitochondrial localization*, or *combined* (in which the partial and mitochondrial outcomes are collapsed) as the outcome levels. Predicted probabilities and 95% confidence intervals were calculated (predict function in R) to compare outcomes.

## Supporting information

a MS-excel file with 9 tabs

## ACKNOWLEDGEMENTS

We thank Dr. Edward Korn for his advice and reagents on *D. discoideum* culturing; Dr. Raúl Covian Garcia for his advice on mitochondria purification; and dictyBase for various plasmids. This work was supported by the Intramural Research Program of National Heart, Lung, and Blood Institute.

## DECLARATION OF INTERESTS

The authors declare no competing interests.

## SUPPLEMENTAL INFORMATION

Supplemental Information includes 9 supplemental tables.

## Supplemental Tables and Table Legends

**Table S1. *D. discoideum* homologs of human mitochondrial proteins**. Homology assessed by BlastP (expect < 0.001 and bit-score > 40) using the Human MitoCarta 3.0 as the query and the *D. discoideum* proteome as the subject. Presented alongside are N-terminal sequence predictions retrieved from TarpetP2 (mTP: mitochondrial targeting peptide; SP: signal peptide; OTHER: neither mTP nor SP).

**Table S2. Proteomics discovery relative enrichments and relative enrichment ratios**. Relative enrichment ratios (columns 2-4) and their respective p-values (columns 5-7) for samples quantified in the proteomic discovery experiment. Relative enrichments from lysate (columns 8 and 9), crude mitochondria (columns 10 and 11), and purified mitochondria (columns 12 and 13) samples. All sample enrichments are calculated from three biological replicates.

**Table S3. Proteomics discovery core mitochondrial proteins**. *D. discoideum* mitochondrial ETC, OXPHOS, and TCA cycle proteins used for enrichment normalization. Information retrieved from uniprot.org.

**Table S4. Proteomics discovery mitochondrial targeting sequence prediction**. N-terminal sequence predictions retrieved from TargetP-2.0 for all samples quantified in the proteomic discovery experiment (n = 6892, columns 1-3) and all proteins with an RER > 0.343 (n = 908, columns 4-6). (mTP: mitochondrial targeting peptide; SP: signal peptide; OTHER: neither mTP nor SP.)

**Table S5. Microscopy validation of relative enrichment ratio cutoff**. Localization patterns of proteins used to verify the RER cutoff. A red highlight indicates proteins that were removed from the final mitochondrial compendium while a green highlight indicates proteins that were added to the final mitochondrial compendium based on the protein’s localization. Proteins that were unable to be synthesized or transfected and proteins that have homology to a protein within the Human MitoCarta 3.0 are indicated in red and blue, respectively.

**Table S6. Classifications of the mitochondrial protein compendium**. Biological process classifications were determined by PANTHER biological process or PANTHER family of the *D. discoideum* protein, or of the *H. sapiens* protein. Homologs of the *D. discoideum* proteins were established by HMMER alone (best homolog) or by HMMER and BlastP (*D. purpureum* whole proteome, *R. prowazekii* whole proteome, *S. cerevisiae* whole proteome and mitoproteome, and *H. sapiens* whole proteome and mitoproteome.

**Table S7. Ribosome proteins within the mitochondrial protein compendium**. List of all mitochondrial proteins with ribosomal protein classifications according to their protein name and/or PANTHER family.

**Table S8. Scaled RNAseq data**. The normalized expression profiles for genes encoding mitochondrial proteins (Parikh *et al*., 2010; Stajdohar *et al*., 2017) scaled to a mean of 0 and standard deviation of 1. (Groups: biological process categories; t0-24: timepoints in hours within the 24-hour starvation-induced development cycle.) Upregulated genes (log2 fold-change ≥ 1) at specific timepoints are indicated in red.

**Table S9. List of oligos used for cloning**. Homology to pDM323 used for ligation is underlined. Reverse primers contain an additional base (bolded) to retain reading frame.

## Notes

### Competing Interest Statement

The authors have declared no competing interest.

